# Structure – silencing duration relationships in RNAi medicines in rapidly dividing cells

**DOI:** 10.1101/2024.09.09.612002

**Authors:** Anastasia Kremer, Tatyana Ryaykenen, Xavier Segarra-Visent, Melanie Sauer, Qi Tang, David A Cooper, Dimas Echeverria, Clothilde Philouze, Emilie Bayon, Dan Georgess, Reka A Haraszti

## Abstract

RNA interference (RNAi)-based medicines offer precise targeting of virtually any transcript, making them an appealing new drug class for addressing unmet needs in immune-oncological applications. While RNAi therapies show exceptional duration of effect in non-dividing cells, their efficacy in rapidly dividing cells, crucial for immune-oncology, remains largely unexplored. Unlike in non-dividing cells, full chemical modification in rapidly dividing cells has not consistently extended silencing duration, according to limited data available.

In this study, we investigated key factors affecting the duration of effect for three main types of RNAi-based therapeutics (siRNA, miRNA mimics, and miRNA inhibitors) in rapidly dividing cancer and immune cells. Saturation of intracellular depots by multiple loading doses, a common strategy to prolong silencing duration in non-dividing hepatocytes, had minimal impact on siRNA duration of effect in rapidly dividing cells. However, modifying the antisense strand with a 5’*-(E)-*vinylphosphonate (5’-VP) to protect siRNAs from exonucleases and enhance AGO2 binding significantly extended siRNA silencing duration to over 30 days both *in vitro* and *in vivo*. For miRNA mimics, extensive stabilization of the antisense strand with phosphorothioates was not effective and led to reduced potency and silencing duration. Interestingly, a shorter duplex region commonly seen in therapeutic siRNAs partially rescued duration of silencing in miRNA mimics with extended phosphorothioate modifications. On the other hand, miRNA inhibitors demonstrated robust reversal of miRNA activity for an impressive 25 days in cancer cell lines.

Our findings enable the rational design of the chemical architecture and administration regimens of RNAi-based therapies in oncology and immunology.

## Introduction

The sustained efficacy of RNA interference (RNAi) drugs is crucial to realize their full therapeutic potential. Over the years, significant advances in delivery systems and metabolic stability have led to a rise in approved short interfering RNA (siRNA) drugs, a key class of RNAi-based therapeutics, with extended durations of action requiring doses only every 1–6 months [1–6]. Some siRNA drugs have demonstrated gene silencing effects lasting up to 680 days after a single administration [7]. However, most clinical-stage siRNAs target non-dividing hepatocytes. Yet, there is growing interest in developing siRNA and other RNAi-based medicines as immune modulators or cancer therapies, focusing on rapidly dividing cells.

Data on siRNA silencing duration in rapidly dividing cells are limited. Inter-study comparison of *in vitro* data show that full chemical modification of siRNAs may not significantly extend silencing (fully modified siRNA silences for 8 days in activated T cells[8] *versus* unmodified siRNAs silences up to 7 days in cancer cell lines[9]). Partial chemical modification shows only marginal improvements in silencing duration *in vitro* [10, 11], though one study demonstrated enhanced siRNA effect duration in primary mouse T cells *in vivo* with partial modification[12].

In general, preclinical data suggest shorter silencing durations in dividing cells compared to non-dividing cells and in *in vitro* compared to *in vivo* studies for both unmodified and chemically modified siRNAs. In rapidly dividing cell lines, unmodified siRNAs’ silencing duration lasts 3–7 days *in vitro* [9, 11], extending to around 10 days in subcutaneous tumors *in vivo* [11]. In non-dividing cells, unmodified siRNAs can maintain silencing for 3-4 times longer than in dividing cells (3–4 weeks both *in vitro* and *in vivo* [11]), while some studies report shorter silencing durations of 9–11 days *in vivo* in liver [12]. Fully modified siRNAs have achieved 8 days of silencing *in vitro* in rapidly dividing activated T cells[8], while 56 days of silencing *in vivo* in murine lungs[13] and 42 days in murine kidneys[14]. However, these organs contain cells with varying proliferation rates, making it unclear which cell types are responsible for the observed silencing durations. In contrast, fully modified siRNAs showed silencing durations of up to 6 months in non-dividing cells in the brain and eyes of mice and non-human primates [13, 15, 16] and up to 680 days in liver of patients[7]. There is no available data comparing the duration of fully modified siRNA effects *in vitro* with their *in vivo* performance in rapidly dividing cells.

Above findings suggest two key points: (i) cell turnover significantly affects siRNA silencing durability, and (ii) chemistries that extend siRNA longevity in non-dividing cells may be insufficient in rapidly dividing cells, necessitating new and innovative chemical strategies. One promising modification is 5’*-(E)-*vinylphosphonate (5’-VP)[17], which has been shown to increase siRNA concentrations in mouse tissues by 2- to 22-fold[14, 18] and enhance mRNA silencing in non-dividing hepatocytes[14] compared to fully modified siRNAs with standard 5’-P or 5’-OH. Potential mechanisms include resistance to 5’-exonucleases[14] and higher binding affinity to AGO2[19]. The use of 5’-VP in siRNAs for rapidly dividing cells has not yet been explored.

During cell division, intracellular siRNA is distributed between daughter cells, leading to a reduction in concentration after each division. This most likely explains why siRNA silencing duration depends on cellular turnover. According to one hypothesis, silencing may last until the intracellular siRNA concentration drops below its IC50. In other words, the excess siRNA present in a cell combined with the cellular turnover rate may mostly explain silencing duration in dividing cells, and metabolic stability may play a minor role in this context.

Available *in vitro* data seem to support this hypothesis, while *in vivo* data is more discrepant: Fully modified siRNAs are present in a 3000-fold excess in hepatocytes than what is loaded to AGO2 [20]. If a similar scenario would be the case in a dividing cell, 3000-fold excess may translate to a silencing duration of about 12 cell divisions. Partially modified siRNAs, with a 50-fold excess (similar to those delivered *via* lipid nanoparticles to liver[21]), may silence for 5–6 cell divisions. This hypothetical calculation aligns with experimentally observed silencing durations of 8 days for fully modified siRNAs[8], and of 3-7 days for unmodified siRNAs[9] in dividing cells, however *in vivo* results are rather discrepant, suggesting that additional mechanisms may play a role.

A non-chemical approach involves repeated dosing to saturate intracellular (endosomal) siRNA depots and all RISCs[20], a strategy used clinically to prolong silencing in non-dividing hepatocytes[22]. This strategy has yet to be tested in rapidly dividing cells.

Another class of RNAi drugs, miRNA mimics, has begun to emerge in preclinical studies in fully modified forms[23–26], following the failure of unmodified versions in clinical trials[27, 28]. Although systematic studies on their effect duration are lacking, miRNA silencing is typically measured 24–72 hours post-transfection *in vitro*[23, 24], and 5[26] to 9[25] days after a single treatment in vivo. Yet another class, miRNA inhibitors, is gaining attention in immuno-oncology, but studies on their silencing duration remain limited, with effects usually measured 1–3 days after treatment[29].

As outlined above, since factors defining the duration of effect of RNAi medicines in rapidly dividing cells seem to differ from those established in non-dividing cells, a comprehensive analysis is needed, which can later guide the rational design of RNAi therapies for immune-oncology applications. In this study, we address these gaps by investigating how 5’-VP and intracellular depot saturation affect siRNA silencing duration in rapidly dividing cancer and immune cells, and how this translates to *in vivo* silencing. We further examine the relationship between IC50 and silencing duration in dividing cells. We provide data on the effect of various stabilization chemistries on the silencing duration of a model miRNA mimic. Finally, we explore the inhibitory duration of a model miRNA inhibitor in rapidly dividing cells.

## Results

To study siRNA silencing durations, we used a previously described fully modified siRNA backbone that combines 2’-F, 2’-OMe, and phosphorothioates[30]. siRNAs were asymmetric, featuring a 16-nt duplex region and a 5-nt fully phosphorothioated overhang on the antisense strand, which facilitates cellular uptake similar to antisense oligonucleotides[31]. To further ensure efficient cellular uptake, all siRNAs were conjugated to cholesterol or myristic acid[32]. A list of the siRNA sequences and modification patterns used can be found in Supplementary Table 1.

### Saturation of intracellular depots do not extend siRNA silencing duration in dividing cells

First, we aimed to investigate the silencing duration in the T cell model line Jurkat using the aforementioned fully chemically modified siRNAs (Fig. 1). We observed silencing durations ranging from 12 days (Fig. 1A, siRNA targeting *WAPAL*, and Fig. 1B, siRNA targeting *AURKA*) to 15 days (Fig. 1C, siRNA targeting *JAK1*[33]). This observed silencing durability corresponds to 6-8 cell divisions (data not shown) and is at least twice as long as that reported for unmodified siRNAs[9, 11] or for fully modified siRNAs[8]. Notably, the observed silencing duration seemed to be independent of IC50 measured in the same cell type (Fig.1. D-F). Specifically, we dosed cells 2- (Fig.1. C.), 90- (Fig.1.A.) and 400-fold (Fig.1.B.) higher siRNA concentration than corresponding IC50s, translating to 1, 6 and 8 cell divisions, respectively, until siRNA concentration drops below IC50. This only partially corresponds to our measurement that Jurkat cells divide 6-8 times over the course of 15 days. Furthermore, although we observed dose-dependent reduction of target mRNA levels, yet, higher doses did not extend the silencing durations (Fig. 1A-C).

**Figure 1.**
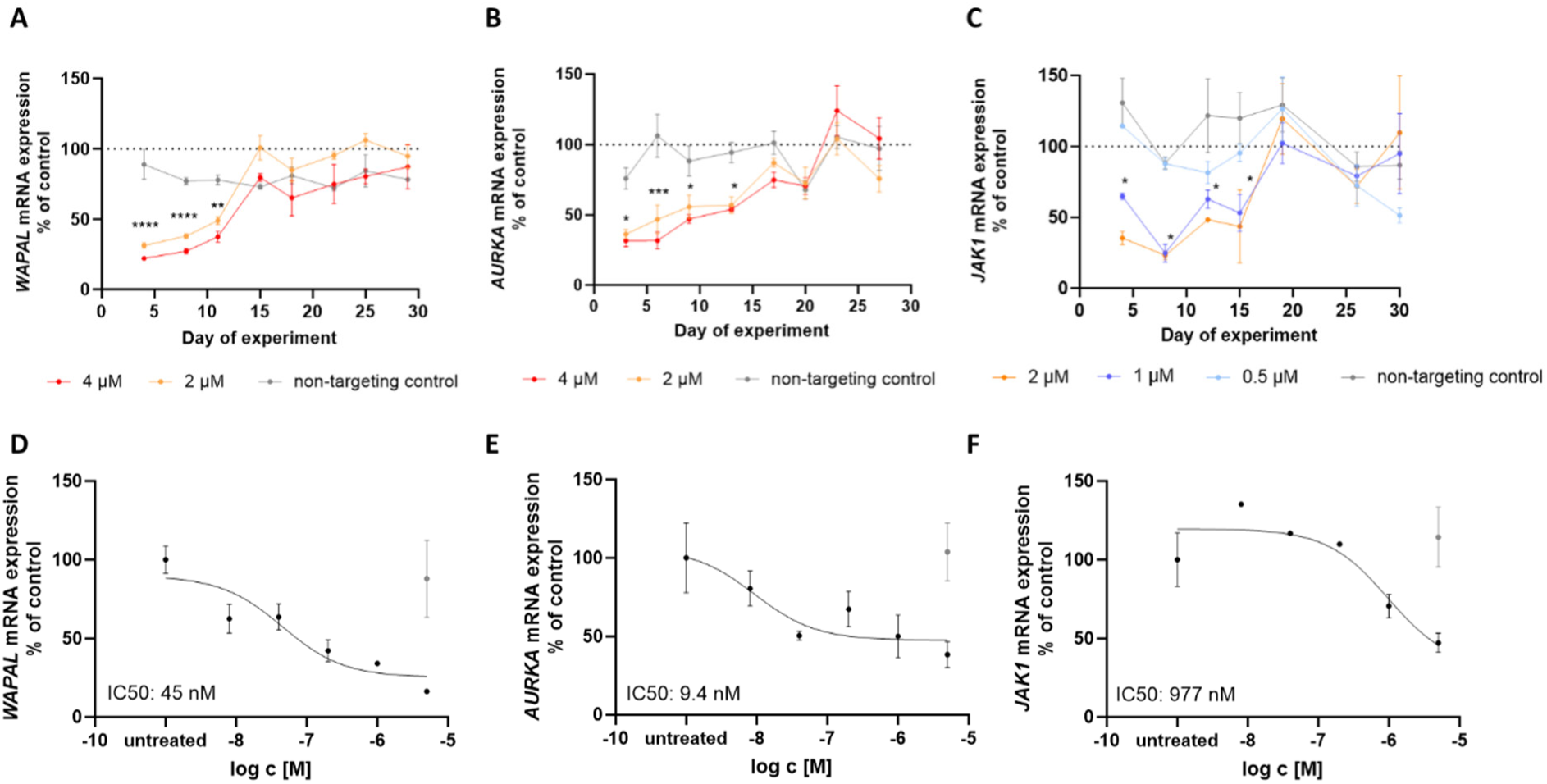
Higher siRNA doses fail to extend duration of silencing in rapidly dividing cells. Jurkat cells were treated with siRNAs targeting *WAPAL* (A and D), *AURKA* (B and E), or *JAK1* (C and F) at the concentrations indicated below the x-axis. Cells were either harvested 6 days post-treatment (D-F) or sampled serially from siRNA-treated cells on the days indicated on the x-axis (A-C). The sampled cells were lysed and frozen. All samples were subjected to QuantiGene Singleplex assays simultaneously for mRNA quantification. IC50 values were determined using the log (inhibitor) – three parameters function in GraphPad Prism. Data points at different time points and treatments were compared using two-way ANOVA with multiple comparison corrections. * p<0.05, **p<0.01, ***p<0.001, ****p<0.0001, N=3-7, mean ± SEM

Initial saturation of intracellular depots with multiple dosing of GalNAc-conjugated siRNAs is a common strategy to sustain potent levels of durable silencing in clinical trials[7]. We aimed to test whether this also applies to rapidly dividing cells *in vitro*. Jurkat cells were treated with three doses of siRNAs, administered three days apart *(i.e.*, on days 1, 4, and 7). This regimen resulted in a silencing duration of 14-21 days (Fig. 2), significantly longer than the 11-day duration observed with a single dose of the same siRNA sequence (Fig. 1.A). However, when silencing duration is calculated from the last treatment (day 7), no additional advantage is observed.

**Figure 2.**
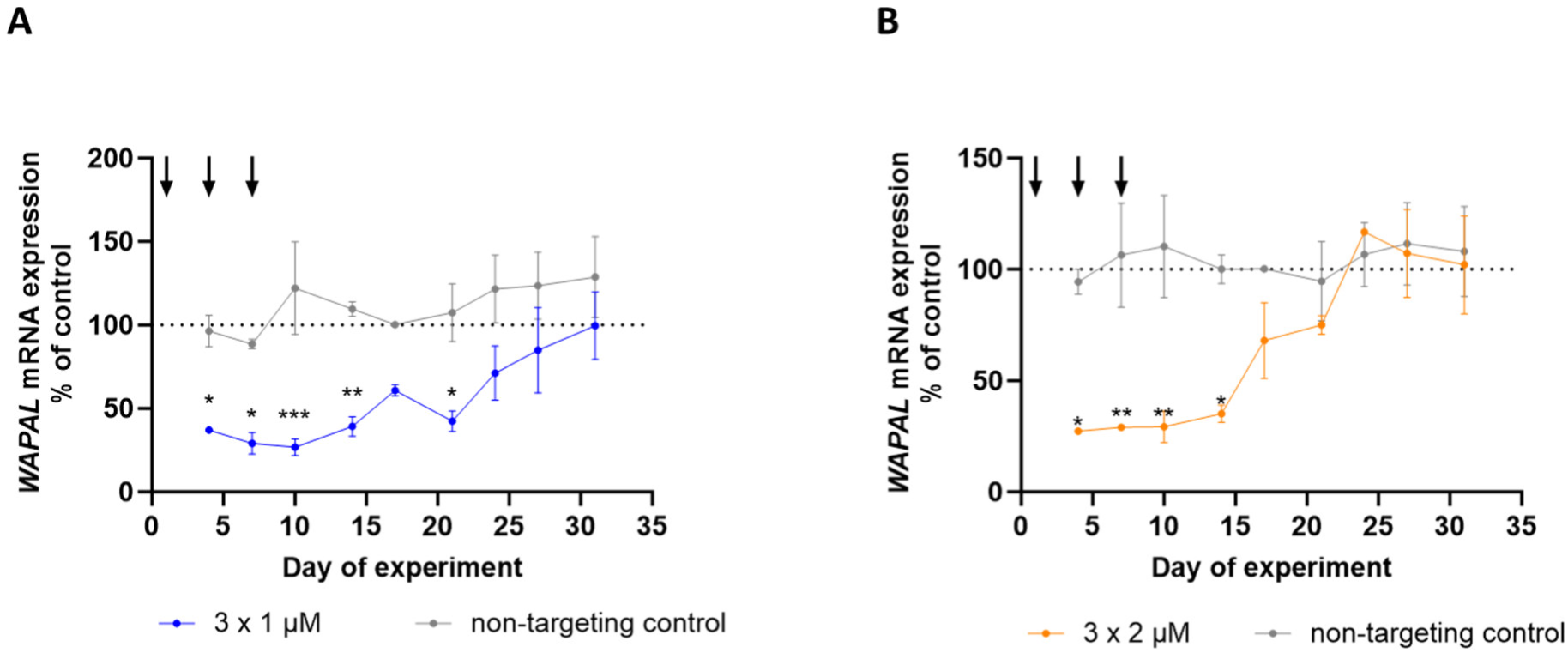
Repeated siRNA dosing fails to extend duration of silencing in rapidly dividing cells. Jurkat cells underwent repeated treatments with siRNAs targeting *WAPAL* at concentrations of either 1 µM (A) or 2 µM (B), up to a maximum of three doses, administered 3 days apart. Cells were harvested from the siRNA-treated cultures on the days indicated on the x-axis, then lysed and frozen. All samples were analyzed simultaneously for mRNA quantity using the QuantiGene Singleplex assay. Data points at different time points and treatments were compared using two-way ANOVA with multiple comparison corrections. * p<0.05, **p<0.01, ***p<0.001, ****p<0.0001, N=3, mean ± SEM

### Silencing duration is independent of cell type

Both immune cells and cancer cells are rapidly dividing and are key targets for RNAi-based therapies. We investigated the silencing duration of the same siRNAs in the adherent cancer cell line HeLa, observing a silencing duration ranging from 7 to 14 days (Fig. 3 A-C), similar to what was observed in Jurkat cells. *WAPAL* silencing decreased faster than *AURKA* or *JAK1* silencing, consistent with the observations in Jurkat cells (Fig. 1). Notably, the silencing duration in HeLa cells remained independent of IC50. Here we treated HeLa cells 22- (Fig.2.C.), 40-(Fig.2.A.) and 100-fold (Fig.2.B.) excess siRNA concentrations than their corresponding IC50s, translating to 4, 5 and 6 cell divisions, respectively, until siRNA concentration drops below IC50. According to our observations, however, HeLa cells have doubling time of roughly 24 hours. Therefore, observed silencing durations of 14-21 days likely correspond to 14-21 cell division events, which is 2-3 times longer than what could be expected from the cell division number needed for critical drop in intracellular siRNA concentration. Hence, in HeLa cells additional mechanisms may play a role in orchestrating siRNA silencing durations compared to Jurkat cells.

**Figure 3.**
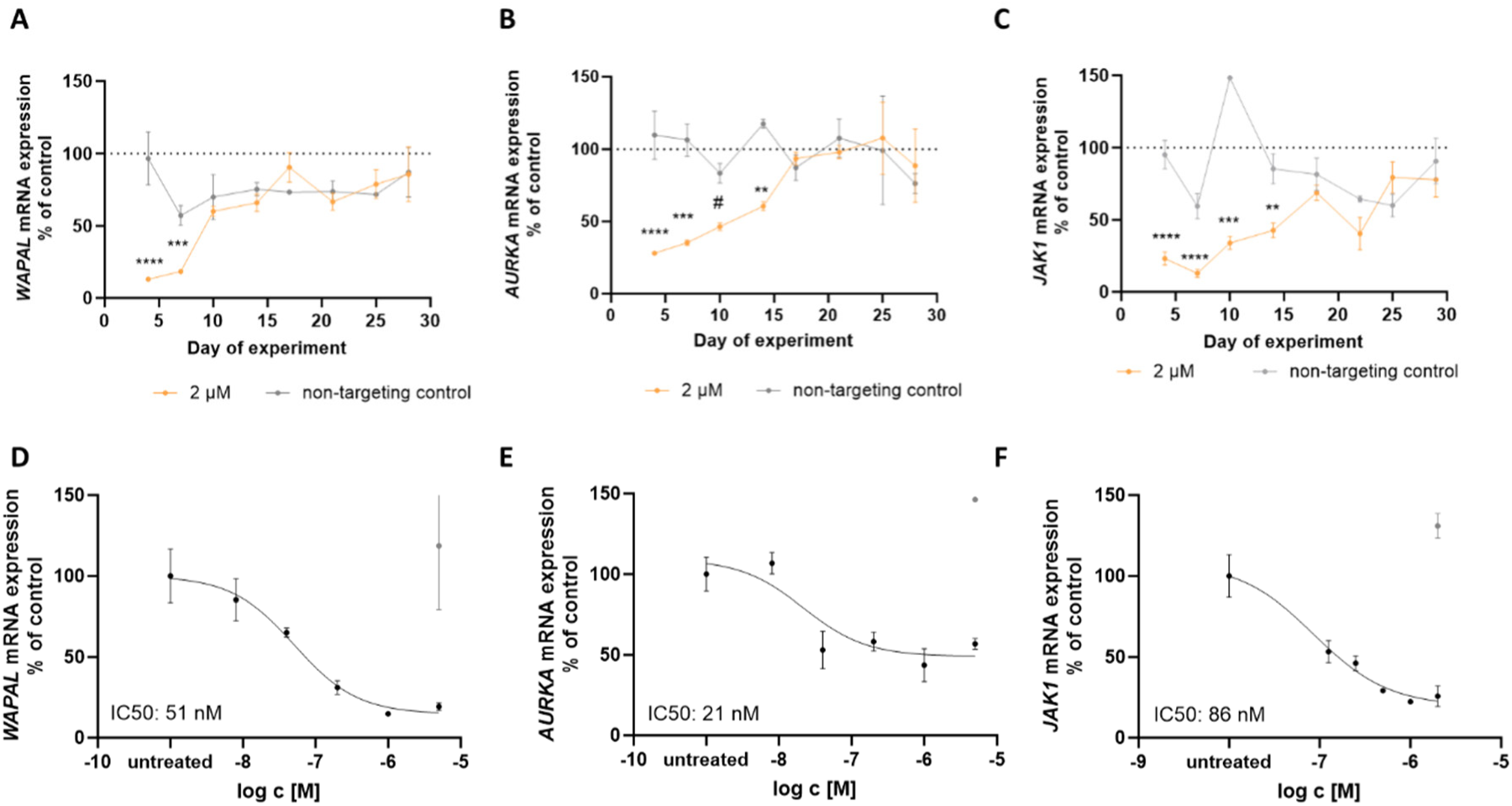
Durations of silencing is independent of cell type and IC50 in rapidly dividing cells. HeLa cells were exposed to siRNAs targeting *WAPAL* (A and D), *AURKA* (B and E), or *JAK1* (C and F) at the concentrations shown below the x-axis. Cells were either collected 6 days after treatment (D-F) or sampled periodically from siRNA-treated cells on the days indicated on the x-axis (A-C). The collected cells were lysed and frozen. All samples underwent simultaneous mRNA quantification using QuantiGene Singleplex assays. IC50 values were calculated using the log (inhibitor) – three parameters function in GraphPad Prism. Data points from various time points and treatments were compared using two-way ANOVA with multiple comparison corrections. * p<0.05, **p<0.01, ***p<0.001, ****p<0.0001, N=3-9, mean ± SEM

We next tested silencing durations in primary activated T cells, a key rapidly proliferating target cell type for RNAi-based therapies. Here, we observed silencing durations ranging from 14 to 17 days (Fig. 4. A.-C.), which is slightly longer than in HeLa (Fig.3. A-C.) or Jurkat cells (Fig.1. A-C.), likely due to the slower proliferation rate of primary T cells compared to HeLa and Jurkat. Indeed we found the doubling time of activated T cells to be about 90 hours versus 24 hours in Hela and 41 hours with Jurkats. Interestingly, we found the potency (estimated based on IC50, Fig. 4 D-F) of the same siRNAs to range from 16 times lower in activated T cells than in HeLa (Fig. 3E versus Fig. 4E) and 37 times lower to 3 times higher than in Jurkat (Fig. 1E versus Fig.4E), yet the duration of silencing remained similar. These large differences in IC50 between cell types may be partially attributed to different siRNA uptake efficiencies, different intracellular siRNA trafficking, and the biological role of the target mRNA in the tested cell types as well as potentially the target mRNA turnover or availability to RNAi, which may also depend on the cell type.

**Figure 4.**
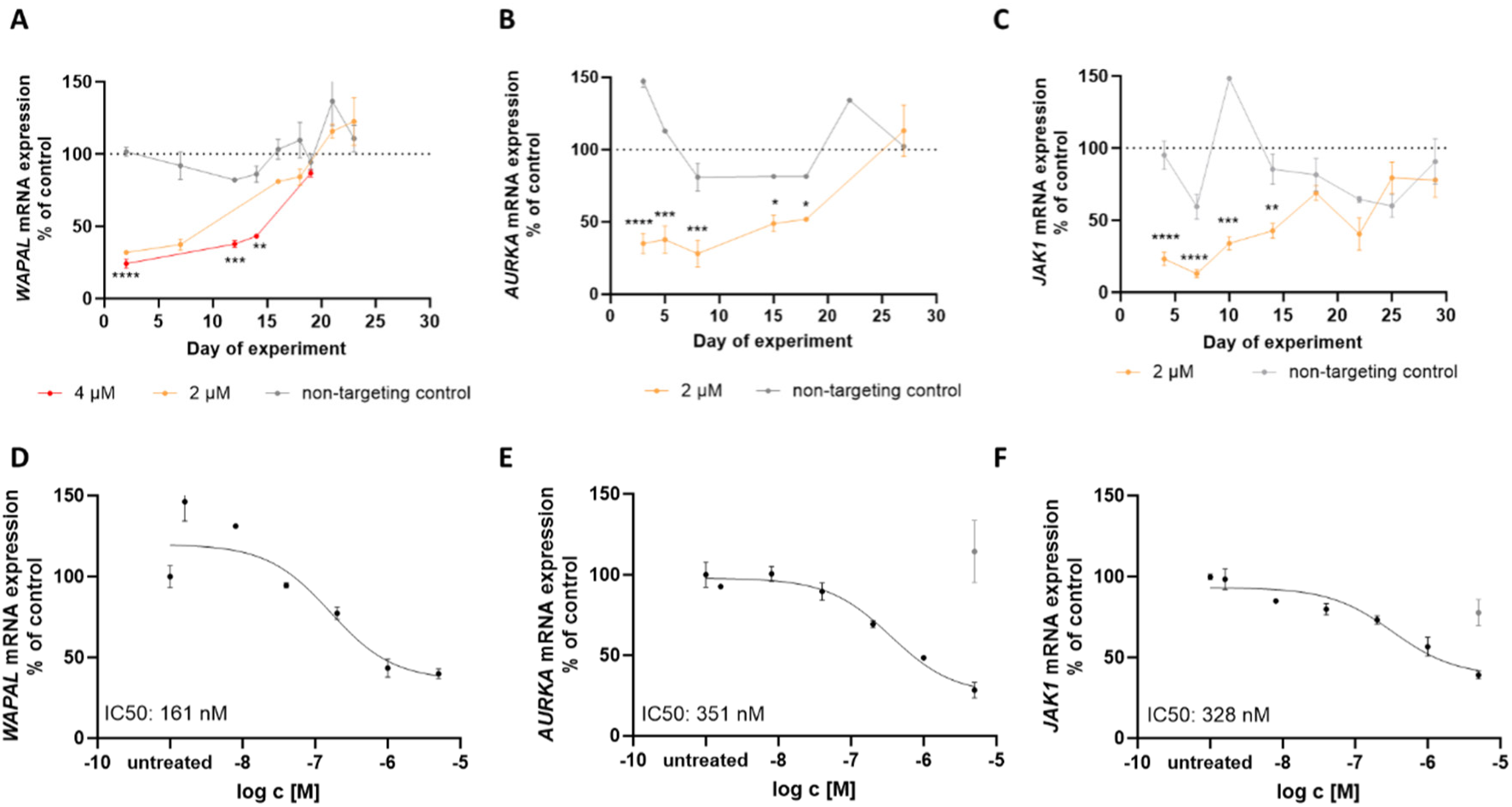
Fully chemically modified siRNAs support biologically relevant silencing durations in primary T cells. Primary human T cells were treated with siRNAs targeting *WAPAL* (A and D), *AURKA* (B and E), or *JAK1* (C and F) at the concentrations indicated below the x-axis. Cells were either harvested 6 days post-treatment (D-F) or sampled at intervals from siRNA-treated cells on the days indicated on the x-axis (A-C). The harvested cells were lysed and frozen. All samples were simultaneously analyzed for mRNA levels using QuantiGene Singleplex assays. IC50 values were determined using the log (inhibitor) – three parameters function in GraphPad Prism. Data points from different time points and treatments were compared using two-way ANOVA with multiple comparison corrections. * p<0.05, **p<0.01, ***p<0.001, ****p<0.0001, N=3-7, mean ± SEM

### 5’-(*E*)-vinylphosphonate stabilization enhances silencing duration

Stabilization of the antisense strand 5’ end *via* 5’-VP has been shown to be beneficial in *in vivo* siRNA applications, enabling silencing in extrahepatic tissues and prolonging silencing duration in kidneys and liver[14]. We treated the model T cell line Jurkat with 5’-P and 5’-VP-siRNAs of two different sequences and at two different concentrations (Fig. 5). We observed a significantly extended silencing duration with both siRNAs when using 5’-VP stabilization of the antisense strand (Fig. 5.A and Fig. 5.D).

**Figure 5.**
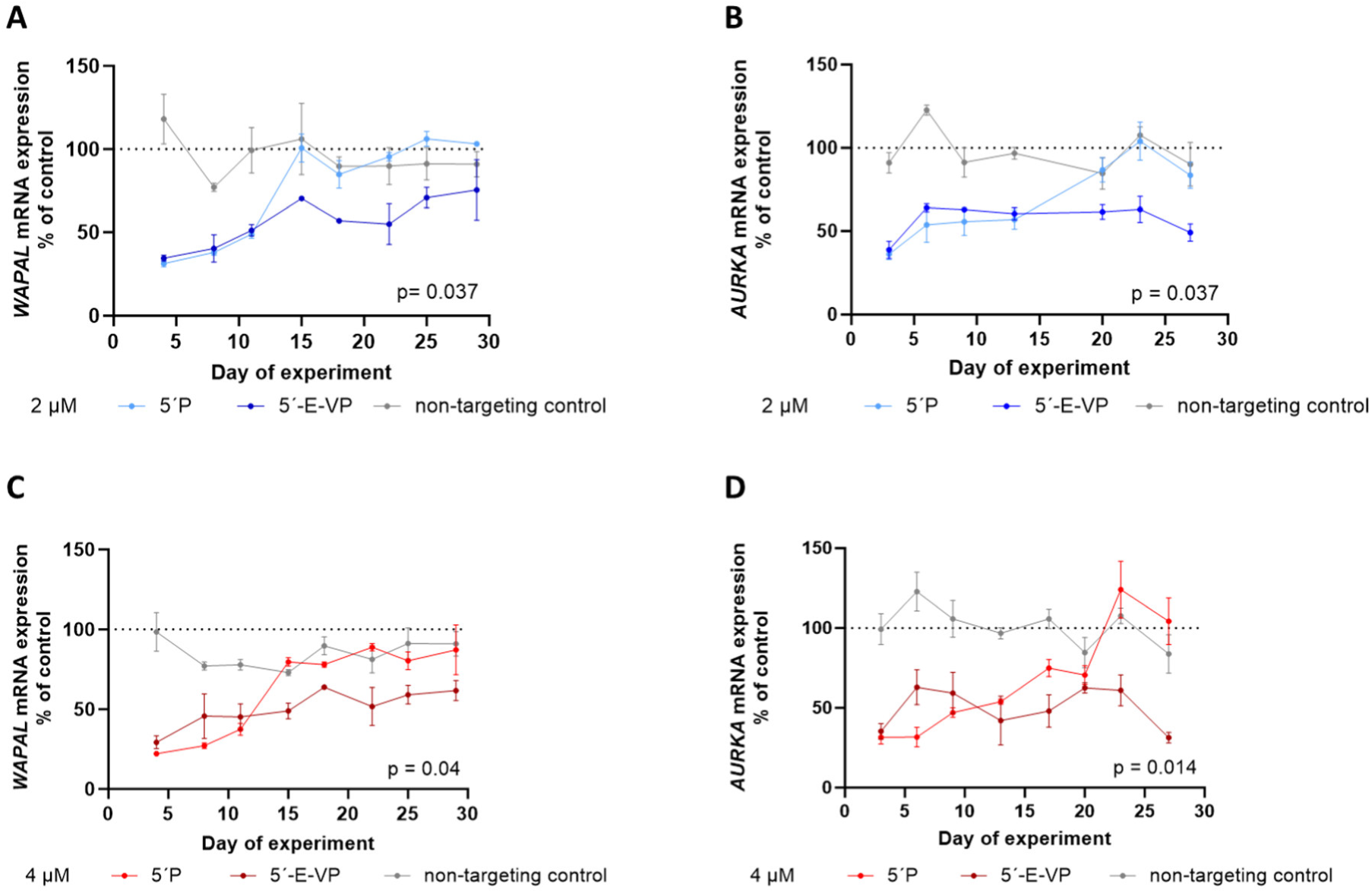
5’-(*E*)-vinylphosphonate stabilization substantially extends silencing durations in rapidly dividing cells. Jurkat cells were treated with siRNAs that were either simply phosphorylated or stabilized with 5’-VP at the 5’ end of the antisense strand. The siRNAs targeted *WAPAL* (A and C) or *AURKA* (B and D) at concentrations of 2 µM (A-B) or 4 µM (C-D). Samples were periodically collected from the treated cultures as indicated on the x-axis. These samples were frozen and later thawed simultaneously for mRNA quantification using QuantiGene Singleplex assays. Data points from various time points and treatments were compared using two-way ANOVA with multiple comparison corrections. N=3, mean ± SEM

We then tested the dose-dependence of silencing duration using a previously identified model siRNA targeting *PPIB*[32, 34, 35], which was stabilized with 5’-VP. Unlike in our previous experiments using 5’-P-siRNAs (Fig.1-4.) silencing duration clearly depended on the siRNA dosing when using 5’-VP, with higher concentrations leading to longer silencing durations (Fig.6.A-B.). At a concentration of 2 µM, 5’-VP-siRNA maintained silencing for beyond 30 days in Jurkat cells (Fig.6. A-B.), corresponding to at least 18 cell division events (data not shown). These findings collectively suggest that there may be an exonuclease-rich environment in dividing cells that can be counteracted *via* the use of 5’-VP stabilization.

**Figure 6.**
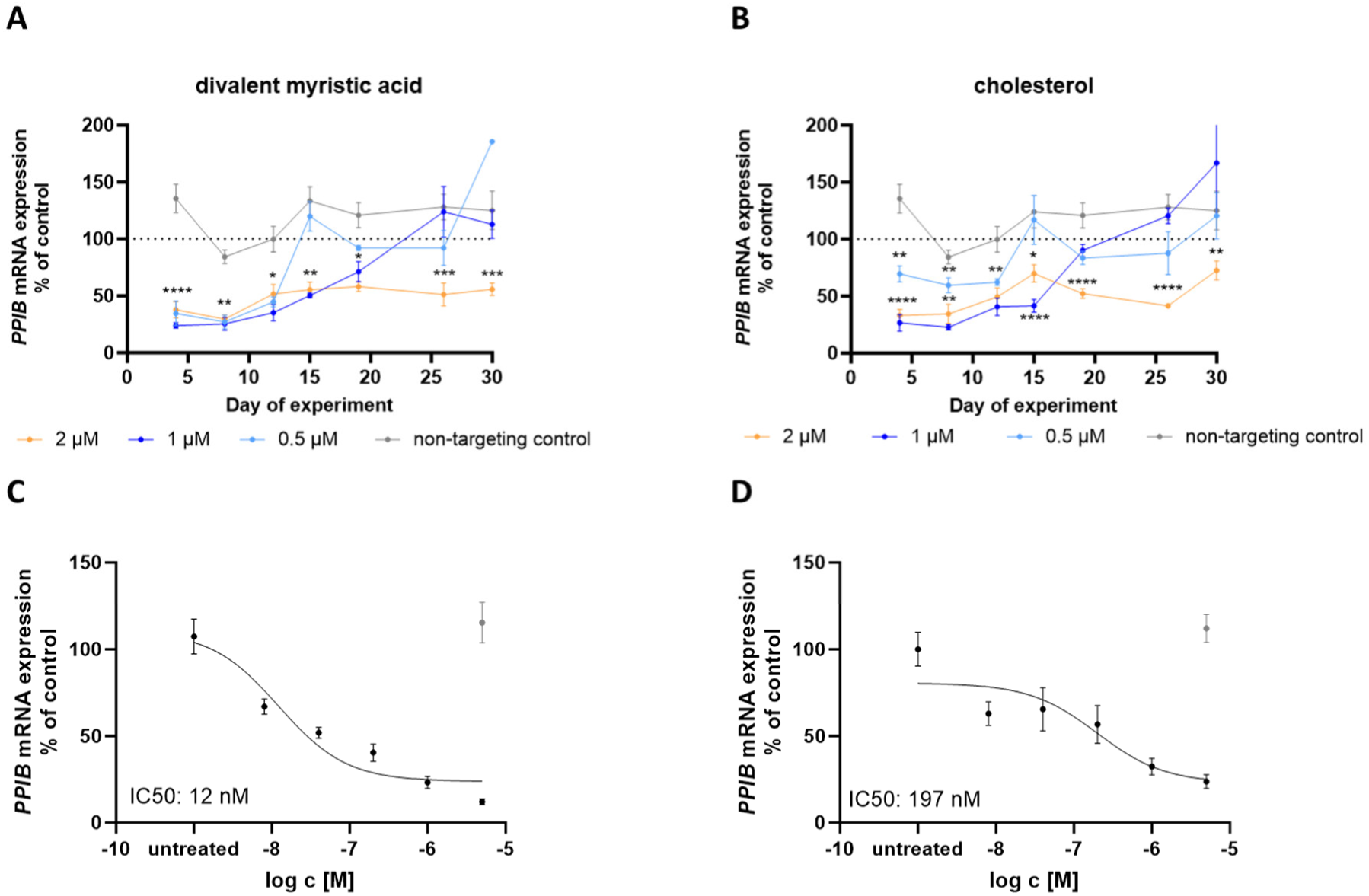
5’-(*E*)-vinylphosphonate containing siRNAs support silencing duration beyond 30 days ex vivo in rapidly dividing cells. Jurkat cells were treated with siRNAs stabilized with 5’-VP and targeting *PPIB* at various concentrations as indicated in the color code (2 µM in orange, 1 µM in dark blue, 0.5 µM in light blue, A-B) or on the x-axis (C-D). The siRNAs were covalently conjugated to either a divalent myristic acid moiety (A and C) or cholesterol (B and D). Cells were either harvested 6 days post-treatment (B and D) or sampled periodically as indicated on the x-axis (A and C). Samples were frozen, later thawed, and mRNA was quantified using QuantiGene Singleplex assays. Data points from various time points and treatments were compared using two-way ANOVA with multiple comparison corrections. * p<0.05, **p<0.01, ***p<0.001, ****p<0.0001, N=3-6, mean ± SEM

The dose-dependence and the length of silencing duration were similar in cholesterol-conjugated (Fig.6.B) and divalent-myristic-acid-conjugated (Fig.6.A) 5’-VP-siRNAs, even though the type of lipid conjugate likely affected the cellular uptake mechanism and therefore the IC50 of these compounds (Fig. 6. C-D.). This finding further supports the notion that silencing duration can be independent of a compound’s silencing potency.

Next, we tested 5’-VP stabilized siRNA in primary activated T cells, and observed a silencing duration beyond 27 days (Supp. Fig. 2.), which corresponded to 8 cell divisions. This silencing duration is notably longer than what we and others observed with 5’-P-siRNAs previously (12-15 days, Fig.4. and 8 days[8]) in activated T cells. Yet, duration of effect in primary activated T cells was notably shorter when measured in the number of cell divisions than what we observed in cell lines.

### 5’-(*E*)-vinylphosphonate stabilization supports sustained silencing durations in immune cells *in vivo*

Since *in vivo* silencing durations of siRNAs have been reported to be longer than *in vitro,* we set out study siRNA silencing duration *in vivo*. For these studies we chose to use 5’-VP-siRNAs, which showed the longest silencing durations *in vitro*.

siRNAs are proposed as modifiers for various cell therapies[36–38]. A significant concern is whether their effect duration is sufficient to induce a meaningful phenotype change once the cell therapy is administered and the cells begin to divide rapidly[39]. To model this scenario, we injected human peripheral blood mononuclear cells (PBMCs) into immunodeficient mice. In this setting, human PBMCs rapidly divide and repopulate the mouse’s immune system, creating a humanized mouse model. We co-incubated PBMCs with a cocktail of three 5’-VP-stabilized siRNAs for 24 hours, then removed the siRNAs and injected the treated cells into the mice.

At 35 days post-injection, the number of human cells usually plateaus at approximately 10 000-fold expansion, corresponding to around 13 cell division events. This level of expansion is comparable to what has been observed with CAR-T cell expansion in patients within the first month of administration[40]. Remarkably, we observed significant silencing of all three siRNA targets at this time point (Fig. 7. A-C), with effect sizes ranging from 83% to 64%.

**Figure 7.**
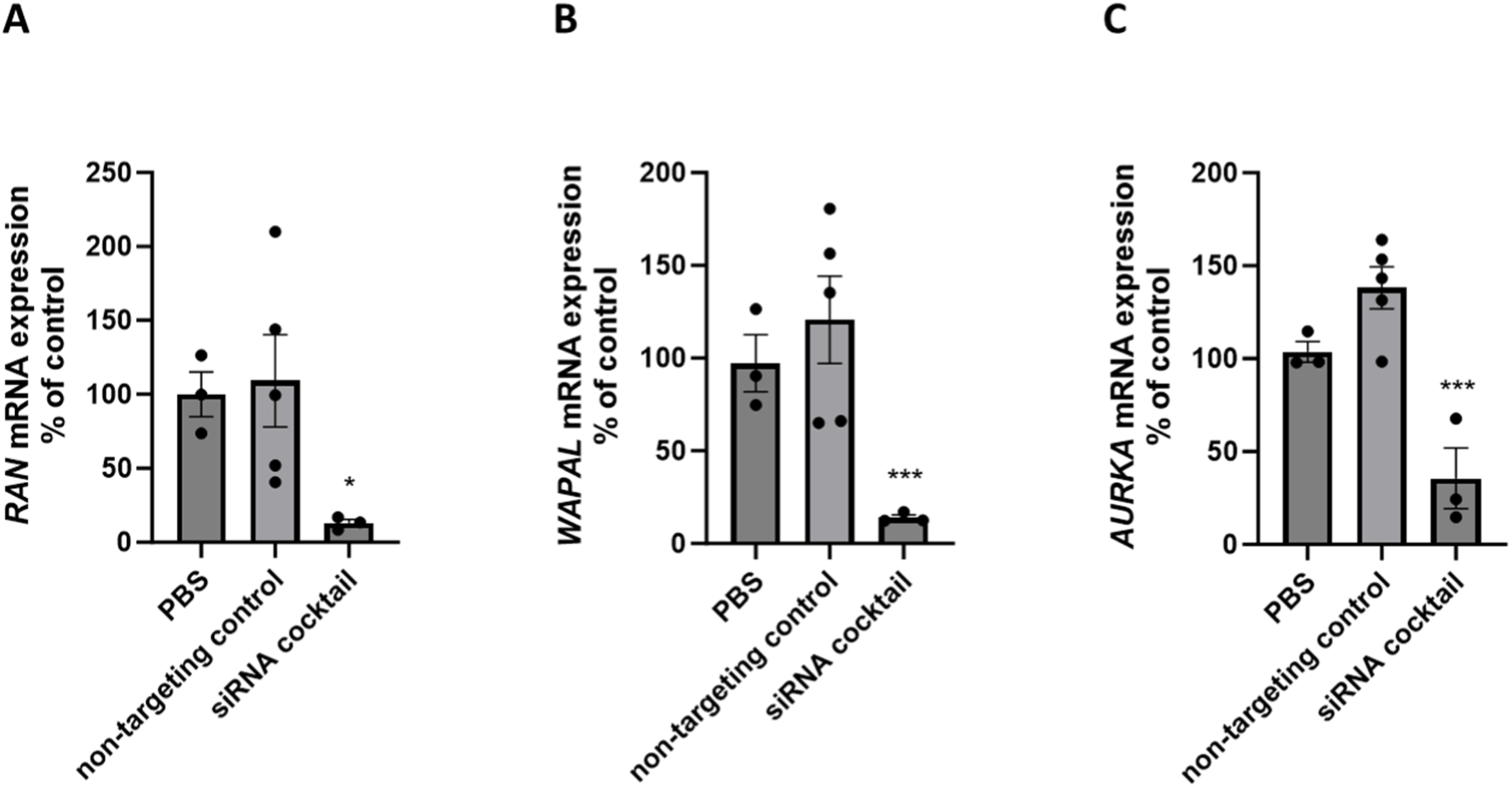
5’-(*E*)-vinylphosphonate containing siRNAs support silencing duration beyond 30 days in vivo in human leukocytes. Human peripheral blood mononuclear cells (PBMCs) were co-incubated with an siRNA cocktail containing three components for 24 hours. The cocktail included siRNAs targeting human *WAPAL*, *AURKA*, and *RAN*, each at a concentration of 3 µM, or a 9 µM non-targeting control siRNA. Following incubation, the siRNA was removed, and the PBMCs were concentrated *via* centrifugation before being intraperitoneally injected into immunodeficient NCG mice. Mice were harvested 33 days post-treatment, and human *WAPALl*, *AURKA*, and *RAN* mRNA levels were quantified in whole blood. Data were analyzed using one-way ANOVA with multiple comparison corrections. * p<0.05, **p<0.01, ***p<0.001, ****p<0.0001, N=3-5, mean ± SEM

### miRNA mimic phosphorothioate backbone interferes with duplex structure to define silencing duration

We next investigated the silencing duration in a model miRNA mimic, miR-146a, using a previously published miRNA mimic chemical scaffold[23] in a dual fluorescent reporter system[41] in HeLa cells. The miRNA mimic had the sequence of mature human miR-146a antisense and sense strand (miRbase[42]). The miRNA mimic was fully chemically modified with an alternating pattern of 2’-F and 2’-OMe and two phosphorothioate (PS) linkages on each end of both strands[23]. We observed a striking 21-day-long silencing duration of this compound in a fluorescent reporter assay in HeLa cells (Fig.8.A., blue). This is notably longer than the silencing durations observed with siRNAs (Fig. 1, 3, and 4). To determine whether these differences could be due to measuring mRNA silencing for siRNAs *versus* protein silencing for the miRNA mimic, we tested protein silencing with siRNAs using a reporter luciferase assay. We observed luciferase silencing for up to 21 days (Supp. Fig. 2), which was comparable to the duration observed with the miRNA mimics. Since siRNA action often results in greater protein silencing than mRNA silencing[15], this may explain the longer effect duration observed when assessing protein levels compared to mRNA levels.

**Figure 8.**
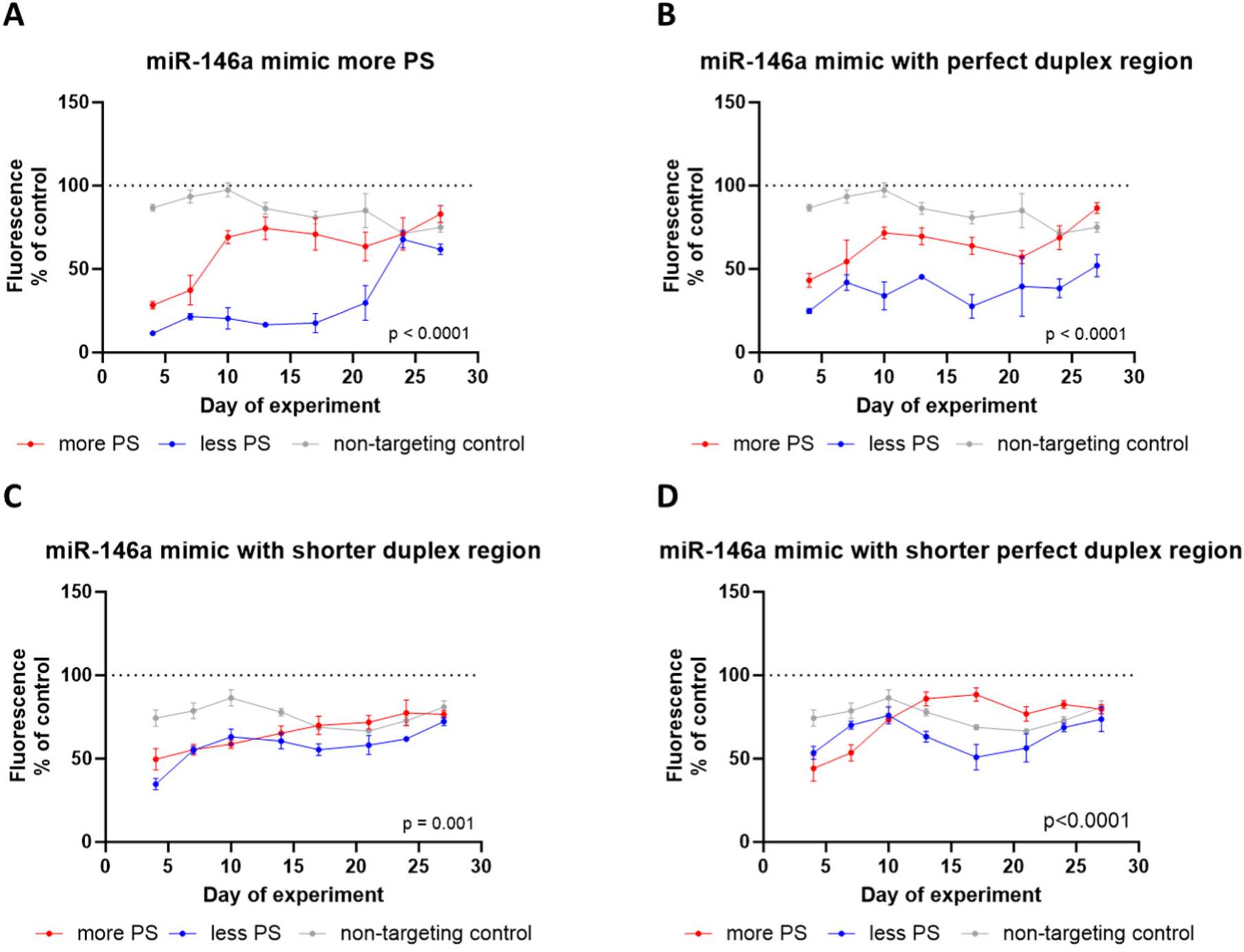
miRNA mimic phosphorothioate content, duplex length and duplex complementarity affects duration of silencing in rapidly dividing cells. HeLa cells were initially transduced with a dual fluorescent reporter system. This system expressed a miR146a complementary sequence with a 4-nucleotide central bulge mismatch in the 3’ UTR of ZsGreen[41]. DsRed was used as a control for transduction efficiency. The transduced HeLa cells were then treated with fully chemically modified miR146a mimics. These treatments included the original human miR146a sequence from miRBase[42] (A), a truncated sense strand of above (C), a fully complementary sense strand (B), or a fully complementary truncated sense strand (D). miRNA mimics sense strand either contained 2 (less PS, blue) or 7 (more PS, red) phosphorothioate (PS) linkages at the 3’ end. Cells were imaged for ZsGreen and Dsred fluorescence using a Tecan Infinite Pro M-Plex plate reader on the days indicated on the x-axis. ZsGreen signal was then normalized to Dsred signal. Data points from various time points and treatments were compared using two-way ANOVA with multiple comparison corrections. N=3-6, mean ± SEM

We next explored, whether the stability of the miRNA mimic duplex structure affected miRNA performance and synthesized a miRNA mimic version with an artificial sense strand fully complementary to the antisense strand, maintaining the same chemical scaffold (Fig.8.B.). Here we observed a slight loss in silencing activity, which in line with reported data[24], yet, the duration of silencing was now maintained for approximately 27 days (Fig.8.B., blue), a slight improvement over the original compound. Next, we aimed to test, whether destabilizing the perfect duplex region may rescue the slightly reduced silencing activity of the second-generation miRNA mimic. We achieved this destabilization by truncating the sense strand by 5 nucleotides (Fig.8.D.). When using this third generation miRNA mimic, we observed a further reduction in silencing levels, yet silencing duration was still maintained up to 21 days (Fig.8.D., blue). We then asked, whether the failure of the third-generation miRNA mimic to rescue silencing activity of the second generation might have been due to the nuclease-sensitivity of the antisense strand overhang generated by truncated the sense strand. One chemical strategy to overcome this problem come from the siRNA world, where this overhang can by entirely phosphorothioated. Therefore, we tested a fourth generation of miRNA mimic, which now had extended phosphorothioate (PS) stabilization on the antisense strand compared to the third generation (Fig.8.D., red). Unexpectedly, the increase in PS linkages significantly reduced silencing duration of the compound (Fig.8.D., red *versus* blue). To further understand, how the number of PS linkages and the stability of the duplex region affect miRNA mimic silencing duration we tested additional versions, where we either truncated the sense strand in the context of natural miR146a sense strand sequence (Fig.8.C.) or added extended PS linkage to all previous versions (Fig.8. A-D. red). Collectively, we made the following observations: (1) increased number of PS linkages reduced silencing duration in all miRNA mimic versions tested (blue curves showing longer silencing durations than red curves). (2) Truncated sense strand reduced silencing potency but extended silencing duration in the context of natural miR146a sense strand sequence and extended PS linkages (Fig.8.A. *versus* Fig.8.C. red curves). (3) Truncated sense strand only reduced silencing potency in context of less PS linkages but left silencing potency intact in the context of extended PS linkages when used in miRNA mimics with artificial fully complementary sense strand sequence and a perfectly matched duplex region (Fig.8.B. versus Fig.8.D.). In sum, the loss of silencing duration when added extended PS linkages to the original miRNA mimic (Fig.8.A. red *versus* blue) could partially be rescued by truncating the sense strand (Fig.A. red *versus* Fig.8.C.red). Furthermore, miRNA mimic structure with perfectly matched duplex region and 2 P linkages of each end of each strand showed the longest silencing duration (Fig.8.B. blue).

Sequences and chemical modifications patterns of all miRNAs mimics used in this study can be found in Supplementary Table 1.

### Lipid-conjugated miRNA inhibitor de-represses miRNA targets for beyond two weeks

We next evaluated the duration of effect of a modified version of the miRNA inhibitor cobomarsen. Cobomarsen is a single-stranded oligonucleotide composed of DNA and LNA residues with a fully phosphorothioated backbone[29] and an inhibitor miR-155. Cobomarsen is being pursued in clinical trials in various hematological malignancies[29], hence in rapidly dividing cells. We used a cholesterol-conjugated version of cobomarsen (at the 5’ end) in order to enhance cellular uptake.

Initially, we assessed the duration of effect of cholesterol-conjugated cobomarsen in HeLa cell line, which we engineered to constitutively overexpress miR-155 (Fig. 9.A.). Using a dual luciferase reporter with miR-155 target sites in the 3’ UTR, we observed a 2-3-fold increase in luciferase signal, which persisted for 24 days (Fig. 9.A.). Subsequently, we tested the effect duration of modified cobomarsen in the Jurkat cell line, which endogenously expresses miR-155. In this case, we observed a 50% increase of the validated miR-155 target, *MYD88* expression, which lasted for 25 days (Fig. 9.B.). The fold change in de-repression of miR-155 target was consistent with previous reports[29], despite using a 2 to 5-fold lower concentration of the miR-inhibitor[29]. In general, we found a larger de-repression of miR-155 targets in miR-155-expressing HeLa than in Jurkat. This observation may be due to most likely higher expression levels of miR-155 in lentivirally transduced HeLa than in wild-type Jurkats.

**Figure 9.**
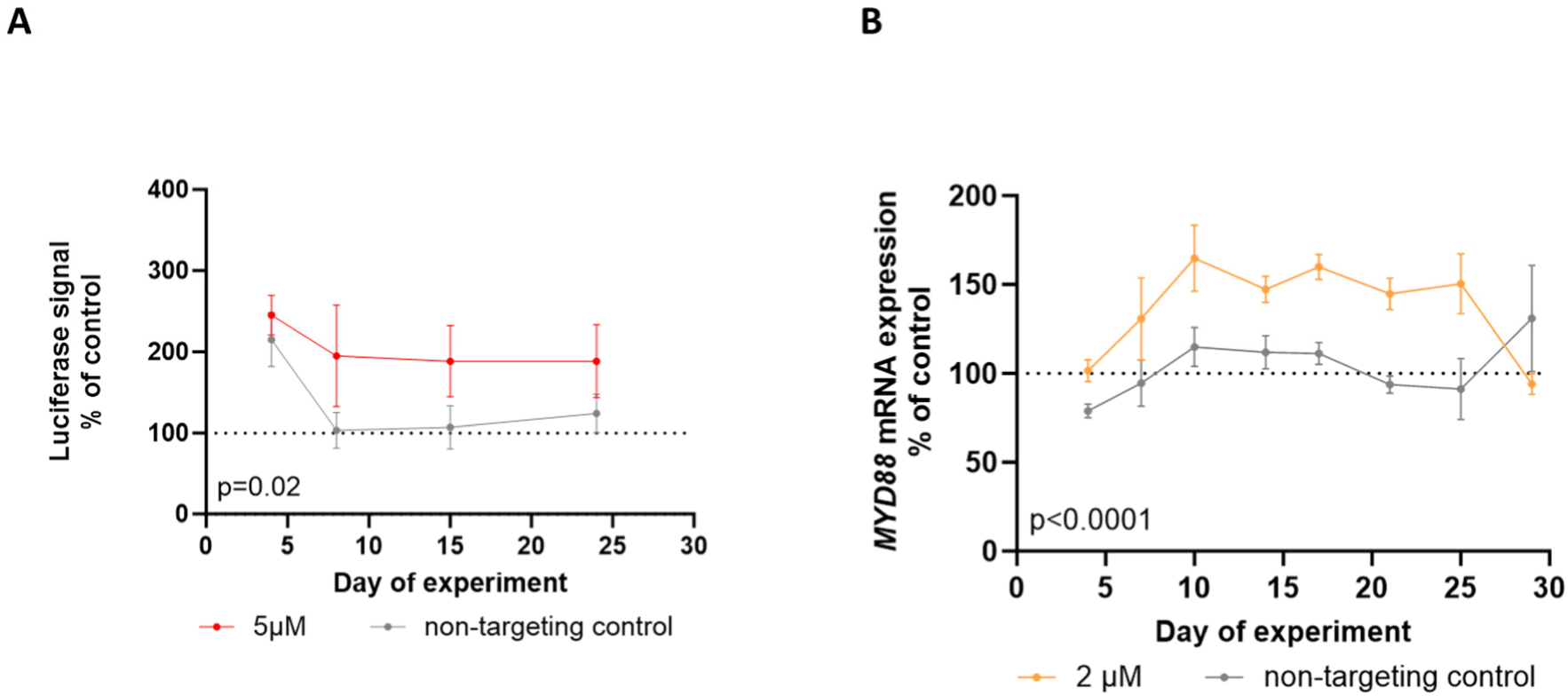
miR-155 inhibitor shows duration of silencing beyond three weeks in rapidly dividing cells. HeLa cells were initially transduced with a miR-155-expressing lentivirus and then transfected with a DualGlo plasmid containing the miR-155 target sequence cloned four times in tandem into the 3’ UTR of Rluc. The transfection was repeated once a week. The transduced and transfected cells were treated with a cholesterol-conjugated miR-155 inhibitor (A) and luminescence was measured using the DualGlo assay. The Rluc signal was normalized to the Fluc signal. Jurkat cells were also treated with the cholesterol-conjugated miR-155 inhibitor, and the miR-155 target, *MYD88* mRNA, was quantified using the QuantiGene Singleplex assay on the days indicated on the x-axis (B). Data curves from various time points and treatments were compared using two-way ANOVA. N=3-10, mean ± SEM

## Discussion

In this study, we provide the first systematic analysis of the duration of effects for various RNAi-based therapies in rapidly dividing cell types, with a focus on fully chemically modified oligonucleotides. For all three classes of RNAi drugs—siRNA, miRNA mimics, and miRNA inhibitors—we show that full chemical modifications enable sustained effects lasting at least 3 weeks in the cancer- and immune cells tested. These findings alleviate concerns about the ability of RNAi-based medicines to achieve clinically meaningful effect durations in immune-oncological applications. Specifically, we show that siRNA silencing duration lasts well beyond 35 days and 13 cell division events in therapeutic cells injected *in vivo*. Such effect durations are able to support therapeutic dosing regimens that are compatible with current immuno-oncology drug administration schedules. Furthermore, the effect durations reported here are likely to cover the entire biologically relevant lifespan of human T cells, which range from 7 to 15 cell division events[43].

Mechanistically, we found that stabilizing siRNA antisense strands with 5’-VP meaningfully extends silencing duration. 5’-VP siRNA has not shown improvement in short-term *in vitro* assays over 5’-P siRNA in previous reports[14], which is in line with our current findings. Yet, 5’-VP siRNAs have shown up to 22-fold enhancement in tissue accumulations in the short term *in vivo*, translating to increased silencing levels[14]. Here, we are first to show that 5’-VP stabilization of siRNA sense strands is crucial for applications in dividing cells and is able to extend silencing duration without increasing the level of silencing. Collectively, these observations suggest that dividing cells may present an intracellular environment potentially rich in exonucleases, where exonuclease resistance *via* 5’-VP[14] is beneficial. Additionally, the balance of kinases and phosphatases is known to be altered during mitosis[44], which in turn may influence the phosphorylation status of fully chemically modified siRNAs. In this context a stable phosphate analogue, such as 5’-VP may be beneficial. 5’-VP has also been shown to enhance siRNA binding *via* AGO2[18]. In dividing cells, it may more frequently be necessary to replenish the available AGO2 pool with new siRNA released from endosomes, and enhancing AGO2-binding of siRNAs *via* 5’-VP may be beneficial in this context.

We found that saturating the hypothesized intracellular siRNA depots with initial multiple dosing failed to significantly prolong silencing duration measured from the last treatment day, unlike reports from clinical trials[7]. This may be explained by the fact that endosomes, which are thought to function as intracellular depots for fully chemically modified siRNAs[45], undergo dynamic changes during mitosis, which may impair their function as siRNA depots.

miRNA mimics tolerated perfectly matched duplexes, however, with reduced silencing potency – in line with existing reports[26]. Extensive PS modification of the 3’ end of the antisense strand was not well tolerated, leading to a moderate loss in silencing activity and profound loss of silencing duration. Size of this effect depended on the length and stability of the miRNA mimic duplex structure and lost silencing duration could partially be mitigated by truncating the sense strand, however, at the cost of further loss in silencing potency. These data indicate that base-pairing outside of the seed region, towards the 3’ end of the sense strand may play a role in silencing duration. Furthermore, our data suggest that potentially different mechanisms may orchestrate silencing potency and silencing duration of miRNA mimics in dividing cells.

miR-inhibitors are in the focus of oncological research, being involved in several clinical trials with dosing regimens ranging from every 3 to 7 days[29, 46]. Our findings suggest that less frequent dosing might also be feasible.

Overall, the data presented here lay the groundwork for the rational design of RNAi-based therapies for immune-oncological applications and of optimized dosing regimens within the context of dividing cells. Notably, we show *in vivo* proof-of-principle that siRNAs may serve as modulators of cell therapies.

## Methods

### Oligonucleotide synthesis, deprotection and purification

Compounds were synthesized using standard solid-phase phosphoramidite chemistry on either a Dr. Oligo 48 high-throughput RNA synthesizer (Biolytic) or using a MerMade 12 (BioAutomation) synthesizer. Standard RNA 2′-*O*-methyl and 2′-fluoro modifications were applied for improving siRNA stability (Chemgenes). The sense strands were synthesized at a 1-μmol scale on a cholesterol-functionalized controlled pore glass (CPG) solid support (Chemgenes) for in vitro experiments. 5-μmol scale was used for in vivo experiments. Custom 5′-(*E*)-vinylphosphonate phosphoramidite (Chemgenes) was applied for *in vivo* studies. For post-synthesis deprotection, sense strands were cleaved from the CPG and deprotected using 40% aqueous methylamine and 30% NH_4_OH (1:1, v/v) at room temperature for 2 h. Guide strands were cleaved and deprotected with 30% NH_4_OH containing 3% diethylamine for 20 h at 35°C. 5′-E-VP containing antisense strands were washed with bromotrimethylsilane:pyridine (3:2, v/v) in dichloromethane while still on solid support previous to deprotection. The deprotected oligonucleotide solutions were filtered to remove CPG residues and dried under vacuum. Compounds for *in vitro* use were precipitated using a modified ethanol precipitation protocol. Compounds for *in vivo* used were HPLC-purified using an Agilent 1290 Infinity II system. Sense strand were purified using a reverse phase column (PRP-C18, Hamilton). Buffer conditions were as follows: eluent A, 50-mM sodium acetate in 5% acetonitrile, and eluent B, 100% acetonitrile. Guide strands were purified using an anion exchange column (Source 15Q, Ge Healthcare). Buffer conditions were as follows: eluent A, 10-mM sodium acetate (pH 7) in 20% acetonitrile, and eluent B, 1 M sodium perchlorate in 20% acetonitrile. Oligonucleotides were detected by measuring peaks with UV absorbance at 260 nm. Peak fractions were automatically collected for confirming identities. The oligonucleotide fractions were quality-controlled by liquid chromatography-mass spectrometry (LC-MS). Desalting was carried out by size exclusion chromatography.

### Cell culture

Jurkat cells were cultivated in RPMI-1640 with stable glutamine and sodium bicarbonate (R2405, SIGMA) supplemented with 10 % FBS (11573397, Fisher Scientific), 1% Penicillin-Streptomycin (P/S) (P0781, SIGMA) and 25 µM HEPES (9157.1, Carl Roth).

Hela cells and HEK-293T cells were grown in RPMI-1640 with stable glutamine and sodium bicarbonate (R2405, SIGMA) supplemented with 10 % FBS (11573397, Fisher Scientific) and 1% Penicillin-Streptomycin (P0781, SIGMA).

All cell lines were cultivated in T25 tissue-culture-treated flasks, fresh medium added 2 times a week. HeLa and HEK-293T cells were passaged once a week 1:10 or at confluency higher than 70%. HeLa cells were detached using 1 ml of accutase (A6964-100ml, SIGMA) according to manufacturer’s instruction. HEK-293T cells were detached by pipetting up and down. Buffy coats from healthy donors were obtained from the Center of Clinical Transfusion Medicine Tuebingen. PBMCs were isolated using density gradient centrifugation. The buffy coat was disinfected with 70% ethanol, transferred to conical tubes, and diluted 1:1 with PBS. The diluted buffy coat was layered over Ficoll (11768538, Fisher Scientific) and centrifuged. The PBMC-containing interphase was collected, washed with PBS, and centrifuged to remove platelets. The pellet was resuspended in RPMI-1640 supplemented with 10% FBS (11573397, Gibco), 1% P/S (P0781, Sigma), 25 mM HEPES (9157.1, Carl Roth) and counted using Neubauer chamber.

### Generation of activated T cells

Bead activated T cells were produced using a T cell activation/expansion kit (130-091-441, Miltenyi Biotec) according to manufacturer’s instructions. Briefly, 1,5 × 10^6^ PBMCs were seeded per well in a 24-well plate (10380932, Fisher Scientific) in RPMI-1640 supplemented with 10% FBS (11573397, Fisher Scientific), 25 mM HEPES (9157.1, Carl Roth) and 1 mM sodium pyruvate (11360070, Gibco). 2 h after seeding, CD2/CD3/CD28 biotinylated beads were added in a 1:2 bead-to-cell ratio and cells were incubated for 2-3 days at 37°C, 5% CO_2_. Cells were collected and medium was changed to above described medium with added 10 ng/ml IL-7 (207-IL, biotechne) and 3 ng/ml IL-15 (247-ILB, biotechne). Activated T cells were passaged every 2-3 days to maintain a concentration of 1,5 × 10^6^ cells/ml per well.

### Cloning and transfection

We designed a gene block containing binding sites for the respective siRNAs, which was then synthesized by Thermo Fisher Scientific (Waltham, Massachusetts). This block was inserted into the 3′ UTR of Rluc in the psiCheck2 plasmid (a generous gift from Dr. Anastasia Khvorova, University of Massachusetts Chan School of Medicine) using XhoI (R0146S, New England Biolabs) and NotI (R3189S, New England Biolabs) restriction enzymes, following the manufacturer’s instructions. Ligation was performed using either Instant Sticky-end Ligase Master Mix (M0370S, New England Biolabs) or T4 DNA Ligase (10339509, Fisher Scientific).

Transfer plasmids for lentiviral transduction were either purchased from Addgene (e.g., miR-146 fluorescent reporter, Addgene #149718) or generated using Gibson Assembly. The lentiviral luciferase reporter was constructed as follows: an insert containing gene block with binding sites for the respective siRNAs, Rluc and Fluc genes was derived from the psiCheck2 plasmid by PCR using the following primers: Fwd: GCGCTGGATCCGTTTAAACGCGGCCGATTCTTCTGACACAACAGTCT and Rev: TTGTAATCCAGAGGTTGATTAGCGATCGCTTACACGGCGATCTTGCC (Microsynth AG, Balgach, Switzerland). The Addgene plasmid #149718 was digested with AsiSI (R0630S, New England Biolabs) and NotI (R3189S, New England Biolabs). A 20 µl reaction containing 0,05 pmol of each fragment and Gibson Assembly Master Mix (17123229, Fisher Scientific) was incubated for 30 min and transformed into TOP10 cells according to the manufacturer’s instructions. The correct clone was confirmed by sequencing.

The miR-155 expressing vector was assembled from Addgene plasmids #149718 and #78126 as follows: miR-155 was first amplified using the following primers: Fwd: CGCTGGATCCGTTTAAACGCGGCCGCTCTGGCTAACTAGAGAACCC and Rev: CGCCGCCAGTCAAGGTCGAGAATTCAGCTGGTTCTTTCCGCCT (Microsynth AG, Balgach, Switzerland). Plasmid #149718 was digested with EcoRI (R3101S, New England Biolabs) and NotI (R3189S, New England Biolabs). A 20 µl reaction containing 0.08 pmol of each fragment and Gibson Assembly Master Mix (17123229, Fisher Scientific) was incubated for 15 min and transformed into NEBStable (C3040I, New England Biolabs) cells according to the manufacturer’s instructions. The correct clone was confirmed by sequencing.

All transfections were performed using Lipofectamine® 2000 (10696343, Fisher Scientific). HeLa cells were detached from the flask, counted, adjusted to the concentration of 0,2 × 10^6^ cells/ml and added 50 µl/10000 cells/well of 96 well flat bottom TC-treated plate. 0,2 µl of Lipofectamin diluted in 4,8 µl of Opti-MEM (11520386, Fisher Scientific) per well in 1,5 ml tube and 100 ng of DNA in 5 µl of Opti-MEM medium per well in 1,5 ml tube. Diluted Lipofectamin and DNA were combined in one tube and incubated for 5 min. Diluted DNA-Lipofectamin were added 10 µl/well and incubated for 37°C 5% CO_2_ 24 hours.

### Lentivirus production and transduction

Packaging psPAX2 (Addgene #12260), envelope pMD2.G (Addgene #12259) and corresponding transfer plasmid were transfected into HEK-293T cells at 70% confluency in 3:1:4 ratio in total amount of 20 mg to T75 flasks using 39 µl of Lipofectamine® 2000 (10696343, Fisher Scientific). After 24 h of incubation medium was changed. After 48 h virus was harvested and concentrated using 8% PEG8000 (10224963, Fischer Scientific) and 0,14 µM NaCl (10616082, Fischer Scientific).

100 µl of concentrated virus was added to 70% confluent (HeLa) or to 1,0 × 10^6^ cells (Jurkat) in a well of 6 well plate. After 48 h medium was changed. Fluorescence of transduced cells were confirmed using microscopy. Luminescence of transduced cells was confirmed using Dual-Glo Assay (E2920, Promega) according to manufacturer’s protocol.

### mRNA quantification

mRNA was quantified using QuantiGene Singleplex assay. Working Lysis Mixture was freshly prepared before each lysis by adding Proteinase K (QS0106, Life Technologies GmbH) to Lysis Mixture 1:100 (QS0106, Life Technologies GmbH). The Working Lysis Mixture was added to the samples in 1:2 ratio. Then the samples were mixed and incubated at 55°C for 30 min. Mixed by pipetting once more after incubation and either processed immediately or stored at -80°C until analysis. Samples from -80°C were completely thawed at room temperature and incubated for 15 min at 37°C prior to proceeding with the protocol.

QGS assay (Invitrogen™QuantiGene™ Sample Processing Kit, cultured cells (QS0103, Life Technologies GmbH), Invitrogen™QuantiGene™ Singleplex Assay Kit (QS0016, Life Technologies GmbH)) was performed according to manufacturer’s protocol. HPRT was used as housekeeping gene. The following probes were used: *AURKA* - SA-10135, *HPRT* - SA- 10030, *JAK1* - SA-50455, *MYD88* - SA-50330, *PPIB* - SA-10003, *RAN* - SA-15837, *WAPAL* - SA-3006464.

### Serial sampling in vitro

The cells were seeded at different densities depending on the plate format: 0,5×10^6^ cells/500 µl in 24-well plates, 0,03×10^6^ cells /150 µl in 48-well plates, or 0,01×10^6^ cells/100 µl in 96-well flat-bottom plates. siRNAs were prepared to the required final concentration in Opti-MEM medium (500 µl for 24-well, 150 µl for 48-well, and 100 µl for 96-well formats). Before being added to the cells, the siRNA-Opti-MEM solution was vortexed and briefly centrifuged. In repeated treatment experiments, treated cells were adjusted to 0,5×10^6^ cells per well in 500 µl of medium in a new well, and treatments were performed as described above on days 4 and 7. Each siRNA concentration was assigned to a single well in the 24-well format and to three wells in the 48- and 96-well formats.

The concentration of Jurkat cells was assessed for each well at each sample collection using a Neubauer chamber. Depending on cell concentration, 150-250 µl of cell suspension was transferred to a 1,5 ml centrifuge tube, where cell concentration was adjusted to 0,5×10^6^ cells/ml, then lysed for downstream mRNA quantification or further processed for the Dual-Glo assay. Cells in 24 well format were passaged after each sample collection. 0,5 × 10^6^ cells were transferred to new well and medium volume adjusted to 1 ml. In 96 well format after cells passaged 1:2 once a week after every other sample collection.

HeLa cells were washed with 300 µl of PBS and detached using 70 µl of Accutase at each sample collection. Subsequently, 230 µl of medium was added, and 200 µl of the sample from each well was transferred to 1,5 ml tubes for further analysis (either mRNA quantification or Dual-Glo assay). The remaining 100 µl, representing one-third of the cells, was re-plated for further culture.

Activated T cells were seeded in a 24-well plate (10380932, Fisher Scientific ) with 1,5 ×10^6^ T cells per ml per well. Cells were left to settle for 2 h prior to siRNA treatment. siRNA was added directly to the wells. Each well was counted 3 times a week. 1,5 ×10^6^ cells were left in each well and excess cells were collected in a 1,5 ml microcentrifuge tube (EP0030120086, Eppendorf). Cells were lysed as described in “mRNA quantification”. Lysates were either used immediately for QuantiGene assay or stored at -80°C. Volume of wells was adjusted to 1 ml with RPMI-1640 supplemented with 10% FBS (11573397, Fisher Scientific), 25 mM HEPES (9157.1, Carl Roth) and 1 mM sodium pyruvate (11360070, Gibco).

### Mice

All animal experiments were conducted at Transcure Bioservices, Archamps, France, in accordance with local ethical guidelines. PBMCs were thawed, and the DMSO-containing medium was removed by centrifugation. The cells were resuspended in RPMI-1640 (R2405, Sigma-Aldrich) with 10% FBS (35-076-CV, Corning), 1% P/S (11548876, Gibco), 25 mM HEPES (9157.1, ROTI), and 1 mM Sodium Pyruvate (11360070, Gibco). The PBMC concentration was adjusted to 2×10^6^ cells/ml and cultured in T75 flasks. siRNA was added at 3 µM concentration each (9 µM total), non-targeting siRNA at 9 µM concentration, and the cells were co-incubated with siRNAs for 24 hours. Cells were then counted again, centrifuged, resuspended in PBS. 9 × 10^6^ viable cells were injected intraperitoneally into female NOD-Prkd^cem26Cd52^Il2rg^em26Cd22^ mice. The mice were monitored 2-3 times a week for weight and health condition and euthanized on day 33. EDTA blood was collected *via* intracardiac puncture, stored at 4°C for less than 12 hours, and then lysed for downstream mRNA quantification.

### Data analysis and visualization

Data analysis and visualization were conducted using GraphPad Prism (Version 10.1.1 (323)). Silencing curves were fitted using the “log(inhibitor) vs. response (three parameters)” function. Comparisons of these curves were performed using two-way ANOVA with multiple comparison tests. Silencing data in groups of mice were analyzed using one-way ANOVA with multiple comparison tests.

## Acknowledgement

We thank Anastasia Khvorova for making oligonucleotide synthesis infrastructure available for this project. This work was supported by the German Cancer Aid [70113948 to R.A.H.); and the Faculty of Medicine, University of Tübingen [473-0-0 to R.A.H., 2652-0-0 to R.A.H.]. R.A.H. is further supported by the MINT-Clinician Scientist program of the Medical Faculty Tübingen, funded by the Deutsche Forschungsgemeinschaft (DFG, German Research Foundation) – 493665037.

## Author Contributions

Conceptualization A.K., T.R., R.A.H. Methodology A.K., T.R., R.A.H., D.G. Investigation A.K., T.R., X.S., M.S., Q.T., D.A.C., D.E., E.B., C.P., Writing – Original Draft R.A.H. Visualization A.K., T.R., R.A.H. Supervision R.A.H. Project Administration R.A.H. Funding Acquisition R.A.H.

## References

1. Adams, D., et al., Patisiran, an RNAi Therapeutic, for Hereditary Transthyretin Amyloidosis. New England Journal of Medicine, 2018. 379(1): p. 11–21.

2. Fitzgerald, K., et al., A Highly Durable RNAi Therapeutic Inhibitor of PCSK9. N Engl J Med, 2017. 376(1): p. 41–51.

3. Balwani, M., et al., Phase 3 Trial of RNAi Therapeutic Givosiran for Acute Intermittent Porphyria. New England Journal of Medicine, 2020. 382(24): p. 2289–2301.

4. Adams, D., et al., Efficacy and safety of vutrisiran for patients with hereditary transthyretin-mediated amyloidosis with polyneuropathy: a randomized clinical trial. Amyloid, 2022. 23: p. 1–9.

5. Garrelfs, S.F., et al., Lumasiran, an RNAi Therapeutic for Primary Hyperoxaluria Type 1. New England Journal of Medicine, 2021. 384(13): p. 1216–1226.

6. Goldfarb, D.S., et al., Nedosiran in primary hyperoxaluria subtype 3: results from a phase I, single-dose study (PHYOX4). Urolithiasis, 2023. 51(1): p. 80.

7. Badri, P., et al., Pharmacokinetic and Pharmacodynamic Properties of Cemdisiran, an RNAi Therapeutic Targeting Complement Component 5, in Healthy Subjects and Patients with Paroxysmal Nocturnal Hemoglobinuria. Clin Pharmacokinet, 2021. 60(3): p. 365–378.

8. Ligtenberg, M.A., et al., Self-Delivering RNAi Targeting PD-1 Improves Tumor-Specific T Cell Functionality for Adoptive Cell Therapy of Malignant Melanoma. Mol Ther, 2018. 26(6): p. 1482–1493.

9. Bartlett, D.W. and M.E. Davis, Insights into the kinetics of siRNA-mediated gene silencing from live-cell and live-animal bioluminescent imaging. Nucleic Acids Res, 2006. 34(1): p. 322–33.

10. Bartlett, D.W. and M.E. Davis, Effect of siRNA nuclease stability on the in vitro and in vivo kinetics of siRNA-mediated gene silencing. Biotechnol Bioeng, 2007. 97(4): p. 909–21.

11. Wu, S.Y., et al., 2′-OMe-phosphorodithioate-modified siRNAs show increased loading into the RISC complex and enhanced anti-tumour activity. Nature Communications, 2014. 5(1): p. 3459.

12. Mantei, A., et al., siRNA stabilization prolongs gene knockdown in primary T lymphocytes. European journal of immunology, 2008. 38(9): p. 2616–2625.

13. Brown, K.M., et al., Expanding RNAi therapeutics to extrahepatic tissues with lipophilic conjugates. Nature Biotechnology, 2022. 40(10): p. 1500–1508.

14. Haraszti, R.A., et al., 5΄-Vinylphosphonate improves tissue accumulation and efficacy of conjugated siRNAs in vivo. Nucleic Acids Res, 2017. 45(13): p. 7581–7592.

15. Alterman, J.F., et al., A divalent siRNA chemical scaffold for potent and sustained modulation of gene expression throughout the central nervous system. Nat Biotechnol, 2019. 37(8): p. 884–894.

16. Cheng, S.Y., et al., Single intravitreal administration of a tetravalent siRNA exhibits robust and efficient gene silencing in mouse and pig photoreceptors. Mol Ther Nucleic Acids, 2024. 35(1): p. 102088.

17. Lima, W.F., et al., Single-stranded siRNAs activate RNAi in animals. Cell, 2012. 150(5): p. 883–94.

18. Elkayam, E., et al., siRNA carrying an (E)-vinylphosphonate moiety at the 5΄ end of the guide strand augments gene silencing by enhanced binding to human Argonaute-2. Nucleic Acids Research, 2017. 45(6): p. 3528–3536.

19. Elkayam, E., et al., siRNA carrying an (E)-vinylphosphonate moiety at the 5΄ end of the guide strand augments gene silencing by enhanced binding to human Argonaute-2. Nucleic Acids Res, 2017. 45(8): p. 5008.

20. Nair, J.K., et al., Impact of enhanced metabolic stability on pharmacokinetics and pharmacodynamics of GalNAc-siRNA conjugates. Nucleic Acids Res, 2017. 45(19): p. 10969–10977.

21. Pei, Y., et al., Quantitative evaluation of siRNA delivery in vivo. Rna, 2010. 16(12): p. 2553–63.

22. Gaya, A., et al., Results of a phase 1/2 study of cemdisiran in healthy subjects and patients with paroxysmal nocturnal hemoglobinuria. EJHaem, 2023. 4(3): p. 612–624.

23. Segal, M., et al., Hydrophobically Modified let-7b miRNA Enhances Biodistribution to NSCLC and Downregulates HMGA2 In Vivo. Mol Ther Nucleic Acids, 2020. 19: p. 267–277.

24. Garreau, M., et al., Chemical modification patterns for microRNA therapeutic mimics: a structure-activity relationship (SAR) case-study on miR-200c. Nucleic Acids Res, 2024. 52(6): p. 2792–2807.

25. Zhu, F., et al., MiR-146a alleviates inflammatory bowel disease in mice through systematic regulation of multiple genetic networks. Front Immunol, 2024. 15(1366319): p. 1366319.

26. Abdelaal, A.M., et al., A first-in-class fully modified version of miR-34a with outstanding stability, activity, and anti-tumor efficacy. Oncogene, 2023. 42(40): p. 2985–2999.

27. Hong, D.S., et al., Phase 1 study of MRX34, a liposomal miR-34a mimic, in patients with advanced solid tumours. Br J Cancer, 2020. 122(11): p. 1630–1637.

28. van Zandwijk, N., et al., Safety and activity of microRNA-loaded minicells in patients with recurrent malignant pleural mesothelioma: a first-in-man, phase 1, open-label, dose-escalation study. Lancet Oncol, 2017. 18(10): p. 1386–1396.

29. Anastasiadou, E., et al., Cobomarsen, an Oligonucleotide Inhibitor of miR-155, Slows DLBCL Tumor Cell Growth In Vitro and In Vivo. Clin Cancer Res, 2021. 27(4): p. 1139–1149.

30. Davis, S.M., et al., Chemical optimization of siRNA for safe and efficient silencing of placental sFLT1. Mol Ther Nucleic Acids, 2022. 29: p. 135–149.

31. Ly, S., et al., Single-Stranded Phosphorothioated Regions Enhance Cellular Uptake of Cholesterol-Conjugated siRNA but Not Silencing Efficacy. Mol Ther Nucleic Acids, 2020. 21: p. 991–1005.

32. Biscans, A., et al., Diverse lipid conjugates for functional extra-hepatic siRNA delivery in vivo. Nucleic Acids Res, 2019. 47(3): p. 1082–1096.

33. Tang, Q., et al., Rational design of a JAK1-selective siRNA inhibitor for the modulation of autoimmunity in the skin. Nat Commun, 2023. 14(1): p. 7099.

34. Alterman, J.F., et al., Hydrophobically Modified siRNAs Silence Huntingtin mRNA in Primary Neurons and Mouse Brain. Mol Ther Nucleic Acids, 2015. 4(12): p. e266.

35. Kremer, A., T. Ryaykenen, and R.A. Haraszti, Systematic optimization of siRNA productive uptake into resting and activated T cells ex vivo. Biomed Pharmacother, 2024. 172(116285): p. 116285.

36. Schaible, P., et al., RNA Therapeutics for Improving CAR T-cell Safety and Efficacy. Cancer Res, 2023. 83(3): p. 354–362.

37. Zhang, H., et al., shRNA-mediated gene silencing of HDAC11 empowers CAR-T cells against prostate cancer. Front Immunol, 2024. 15(1369406): p. 1369406.

38. An, N., et al., Anti-Acute Myeloid Leukemia Activity of CD38-CAR-T Cells with PI3Kδ Downregulation. Mol Pharm, 2023. 20(5): p. 2426–2435.

39. Rossi, M., et al., Efficient shRNA-based knockdown of multiple target genes for cell therapy using a chimeric miRNA cluster platform. Mol Ther Nucleic Acids, 2023. 34(102038): p. 102038.

40. Maude, S.L., et al., Chimeric antigen receptor T cells for sustained remissions in leukemia. N Engl J Med, 2014. 371(16): p. 1507–17.

41. Emming, S., et al., A molecular network regulating the proinflammatory phenotype of human memory T lymphocytes. Nat Immunol, 2020. 21(4): p. 388–399.

42. Kozomara, A., M. Birgaoanu, and S. Griffiths-Jones, miRBase: from microRNA sequences to function. Nucleic Acids Res, 2019. 47(D1): p. D155–D162.

43. Obst, R., The Timing of T Cell Priming and Cycling. Frontiers in Immunology, 2015. 6(563).

44. Nasa, I. and A.N. Kettenbach, Coordination of Protein Kinase and Phosphoprotein Phosphatase Activities in Mitosis. Front Cell Dev Biol, 2018. 6(30): p. 30.

45. Brown, C.R., et al., Investigating the pharmacodynamic durability of GalNAc-siRNA conjugates. Nucleic Acids Res, 2020. 48(21): p. 11827–11844.

46. Querfeld, C., et al., Phase 1 Trial of Cobomarsen, an Inhibitor of Mir-155, in Cutaneous T Cell Lymphoma. Blood, 2018. 132(Supplement 1): p. 2903–2903.

