## Supplementary data for "Structure – silencing duration relationships in RNAi medicines in rapidly dividing cells"

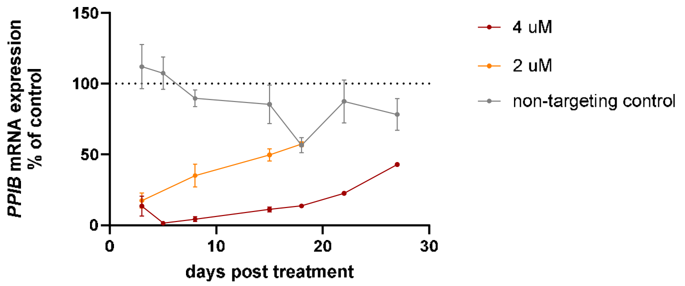


**Supplementary Figure 1. 5'-(*E*)-vinylphosphonate containing siRNA supports sustained silencing in primary T cells ex vivo.** Primary T cells were obtained from human peripheral blood mononuclear cells and polyclonally activated using CD2/CD3/CD28 beads. Cells were then co-incubated with various concentration of siRNAs targeting *PPIB* and mRNA quantified at different timepoints using QuantiGene SinglePlex assay.

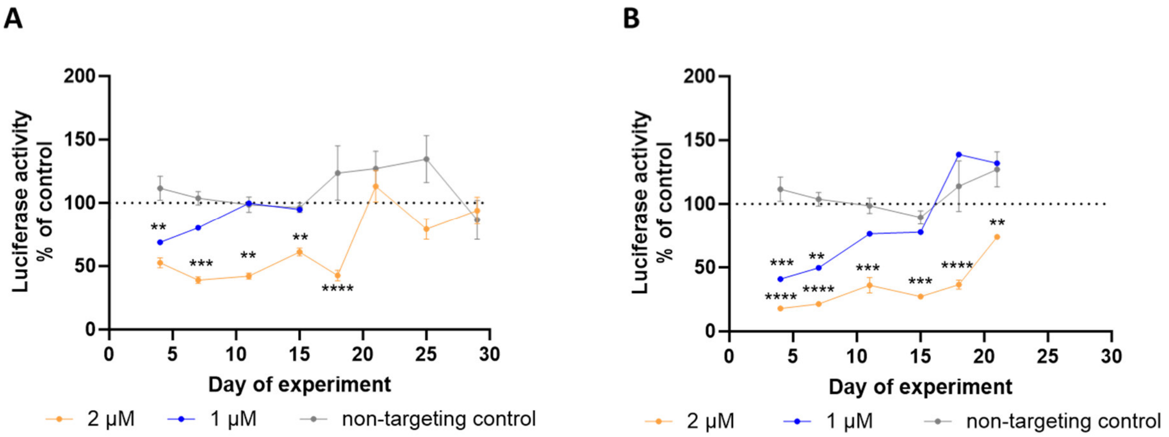


**Supplementary Figure 2. Fully chemically modified siRNAs support sustained protein silencing in reporter assays.** HeLa cells were transduced with a lentiviral dual luciferase reporter, where siRNA target sequence was cloned into the 3' UTR of Renilla Luciferase. Two different siRNAs were used, targeting either *JAK3* (A) or *BIM* (B). Both Renilla and Firefly (*i.e.* trasnduction control) Luciferase activity was measured at different timepoints using DualGlo assay.

### Supplementary Table 1

| Target | Sequence | |
|---|---|---|
| NTC | antisense | P(mU)#(fA)#(mA)(fU)(fC)(fG)(mU)(fA)(mU)(fU)(mU)(fG)(mU)(fC)#(mA)#(fA)#(mU)#(mC)#(mA)#(mU)#(fA) |
| | antisense | V(mU)#(fA)#(mA)(fU)(fC)(fG)(mU)(fA)(mU)(fU)(mU)(fG)(mU)(fC)#(mA)#(fA)#(mU)#(mC)#(mA)#(mU)#(fA) |
| | sense | (mU)#(mU)#(mG)(fA)(mC)(fA)(mA)(fA)(mU)(fA)(mC)(mG)(mA)(fU)#(mU)#(mA)-TegChol |
| *PPIB* | antisense | V(mU)#(fC)#(mA)(fC)(mG)(fA)(mU)(fG)(mG)(fA)(mA)(fU)(mU)#(fU)#(mG)#(fC)#(mU)#(fG)#(mU)#(fU) |
| | sense | (fC)#(mA)#(fA)(mA)(fU)(mU)(fC)(mC)(fA)(mU)(fC)(mG)(fU)#(mG)#(fA)-lipid conjugate |
| *WAPAL* | antisense | P(mU)#(fU)#(mU)(fA)(fG)(fU)(mU)(fA)(mC)(fA)(mU)(fU)(mC)(fU)#(mG)#(fG)#(mU)#(mG)#(mA)#(mG)#(fA) |
| | antisense | V(mU)#(fU)#(mU)(fA)(fG)(fU)(mU)(fA)(mC)(fA)(mU)(fU)(mC)(fU)#(mG)#(fG)#(mU)#(mG)#(mA)#(mG)#(fA) |
| | sense | (mC)#(mC)#(mA)(fG)(mA)(fA)(mU)(fG)(mU)(fA)(mA)(mC)(mU)(fA)#(mA)#(mA)-TegChol |
| *AURKA* | antisense | P(mU)#(fU)#(mG)(fU)(fG)(fU)(mA)(fG)(mC)(fG)(mU)(fU)(mC)(fU)#(mA)#(fG)#(mA)#(mU)#(mU)#(mG)#(fA) |
| | antisense | V(mU)#(fU)#(mG)(fU)(fG)(fU)(mA)(fG)(mC)(fG)(mU)(fU)(mC)(fU)#(mA)#(fG)#(mA)#(mU)#(mU)#(mG)#(fA) |
| | sense | (mC)#(mU)#(mA)(fG)(mA)(fA)(mC)(fG)(mC)(fU)(mA)(mC)(mA)(fC)#(mA)#(mA)-TegChol |
| *JAK1* | antisense | P(mU)#(fC)#(mA)(mG)(mA)(fU)(mU)(mC)(mC)(mU)(mU)(mU)(mG)#(fU)#(mA)#(fC)#(mU)#(mU)#(mC)#(fA) |
| | sense | (mU)#(mA)#(mC)(mA)(fA)(fA)(fG)(mG)(fA)(mA)(mU)(mC)(mU)#(mG)#(mA)-TegChol |
| *RAN* | antisense | V(mU)#(fA)#(mG)(fU)(fU)(fG)(mU)(fA)(mG)(fU)(mU)(fA)(mC)(fU)#(mU)#(fU)#(mU)#(mG)#(mG)#(mC)#(fA) |
| | sense | (mA)#(mA)#(mA)(fG)(mU)(fA)(mA)(fC)(mU)(fA)(mC)(mA)(mA)(fC)#(mU)#(mA)-TegChol |
| *JAK3* | antisense | P(mU)#(fA)#(mU)(fU)(fG)(fA)(mC)(fC)(mC)(fU)(mC)(fU)(mC)(fU)#(mG)#(fU)#(mG)#(mC)#(mA)#(mU)#(fA) |
| | sense | (mA)#(mC)#(mA)(fG)(mA)(fG)(mA)(fG)(mG)(fG)(mU)(mC)(mA)(fA)#(mU)#(mA)-TegChol |
| *BIM* | antisense | P(mU)#(fG)#(mA)(fU)(fC)(fA)(mA)(fC)(mU)(fA)(mA)(fC)(mC)(fA)#(mG)#(fU)#(mU)#(mG)#(mG)#(mG)#(fA) |
| | sense | (mA)#(mC)#(mU)(fG)(mG)(fU)(mU)(fA)(mG)(fU)(mU)(mG)(mA)(fU)#(mC)#(mA)-TegChol |
| miR-146a | antisense less PS | P(mU)#(fG)#(mA)(fG)(mA)(fA)(mC)(fU)(mG)(fA)(mA)(fU)(mU)(fC)(mC)#(fA)#(mU)#(fG)#(mG)#(fG)#(mU)#(fU) |
| | antisense more PS | P(mU)#(fG)#(mA)(fG)(mA)(fA)(mC)(fU)(mG)(fA)(mA)(fU)(mU)(fC)(mC)(fA)(mU)(fG)(mG)(fG)#(mU)#(fU) |
| | sense fully complementary | (mA)#(fA)#(mC)(fC)(mC)(fA)(mU)(fG)(mG)(fA)(mA)(fU)(mU)(fC)(mA)(fG)(mU)(fU)(mC)(fU)#(mC)#(fA)-TegChol |
| | sense truncated | (mC)#(fC)#(mU)(fC)(mU)(fG)(mA)(fA)(mA)(fU)(mU)(fC)(mA)(fG)(mU)#(fU)#(mC)-TegChol |
| | sense fully complementary truncated | (mA)#(fA)#(mC)(fC)(mC)(fA)(mU)(fG)(mG)(fA)(mA)(fU)(mU)(fC)(mA)#(fG)#(mU)-TegChol |
| miR-155-inhibitior | antisense | TegChol-5'(*C)#(**A**)#(*C)#(**G**)#(**A**)#(*T)#(*T)#(**A**)#(*G)#(**C**)#(*A)#(*T)#(*T)#(*A)3' |
| miR-inhibitor NTC | antisense | TegChol-5'(*G)#(**T**)#(*A)#(**G**)#(**G**)#(*A)#(*A)#(**A**)#(*C)#(**A**)#(*C)#(*T)#(*A)#(*C)3' |

| acronym | |
|---|---|
| P | 5‘-phosphate |
| V | 5‘-E-vinylphosphonate |
| # | Phosphorothioate linkage |
| m | 2‘-O-methyl |
| f | 2‘-fluoro |
| * | LNA |
| **bold** | DNA |
| TegChol | Triethyl-glycol-linker conjugated cholesterol |
| NTC | Non-targeting control |

***Supplementary Table 1.*** *Oligonucleotide sequences and their chemical modifications used in this study*
